# Anti-cancer drug Tamoxifen interferes with *Mycobacterium tuberculosis* PhoPR mediated signaling and inhibits mycobacterial growth

**DOI:** 10.1101/2023.12.19.571805

**Authors:** Abhishek Garg, Mansi Pandit, Vandana Malhotra, Deepak Kumar Saini

## Abstract

Two-component signaling (TCS) systems empower all bacteria, including intracellular pathogens like *Mycobacterium tuberculosis (M. tb)* to regulate key pathways governing growth, physiology and virulence. Amongst all *M. tb* TCS systems, PhoPR and DevRS have been studied extensively for their roles in regulating persistence and virulence. Here, we report that besides its cognate response regulator PhoP, the PhoR sensor kinase displays several non-cognate interactions that augment its role in pathogenesis. We demonstrate that PhoR phosphorylates the DevR response regulator and furthermore, is itself subjected to *O*-phosphorylation by PknK, a Ser/Thr protein kinase (STPK), connecting TCS pathways with “eukaryotic-like” STPK driven phosphosignaling. This intersection of non-canonical regulatory pathways and the coregulation of PhoP and DevR regulons make *M. tb* PhoR a potentially attractive drug target. We rationalized that disruption of PhoPR signaling cascade and the resulting dysregulation may result in decreased virulence of *M. tb*. We tested this hypothesis by performing a high-throughput screen for compounds that inhibit autophosphorylation of PhoR sensor kinase. Screening of pharmacologically active, small molecule libraries yielded 11 potential inhibitors, of which one compound, Tamoxifen was able to attenuate PhoR autophosphorylation at micromolar concentrations *in vitro* and *in vivo*. Tamoxifen not only inhibited growth of *Mycobacterium bovis* BCG in culture but also interrupted PhoPR-mediated downstream signaling. Quantitative expression analysis revealed suppression of target gene, *aprA* under acidic conditions. Our findings highlight TCS sensor kinases as promising drug targets and underscore the applicability of clinically relevant anti-cancer drug tamoxifen as a repurposed anti-TB drug.

## Introduction

Tuberculosis (TB) continues to be one of the major causes of death worldwide (1). The causative agent, *Mycobacterium tuberculosis (M. tb),* is a robust pathogenic bacterium that has evolved to adapt and survive within the dynamic environment of the infected host. Despite widespread global efforts, eradication of TB has been a distant dream. The main roadblock has been the persistent or dormant TB bacilli which causes latent tuberculosis infection. It is believed that exposure to the host’s immune responses drive the mycobacteria to undergo transcriptional reprogramming that facilitates its transition into a state of slow growth and non-replicative persistence (2, 3). The fact that mycobacteria can survive in this state for decades provides the impetus to identify novel targets for developing newer therapeutic approaches. Signal transduction systems are central to this adaptability and are known to play a vital role in regulating mycobacterial pathogenesis, persistence, and virulence. One such family of signaling systems is the Two-component signaling (TCS) system which is responsible for sensing environmental cues and modulating the regulation of various transcription factors and gene expression (4, 5). Typically, a TCS system consists of a *sensory component,* usually a Histidine kinase (HK) which senses the environmental stimulus and undergoes autophosphorylation on a conserved Histidine residue. This phosphate is then transferred to a conserved Aspartate residue on a *regulatory component* called the response regulator (RR), a DNA-binding transcription factor capable of regulating the transcription of downstream regulons (6). The other major signaling pathway in mycobacteria is the “eukaryotic-like” Ser/Thr protein kinases (STPKs) (7). The landscape of STPK-mediated *O*-phosphorylation of *M. tb* cellular targets is expansive (8, 9), suggesting an integral role of these proteins in mycobacterial pathobiology.

*M. tuberculosis* has 12 paired two-component systems, six orphan regulators, and two orphan sensor kinases (4, 10). Of these, TCS systems like MtrAB and PrrAB are essential for *M. tb* survival (11, 12), whereas others like PhoPR, and DevRS are implicated in mycobacterial latency and virulence (13–16). The PhoPR TCS system consists of a membrane-bound sensor kinase PhoR that is known to sense low pH and activate a signaling cascade by phosphorylating PhoP response regulator resulting in modulation of its DNA binding property (17, 18). The PhoPR regulon is known to be responsible for regulating the synthesis of complex cell wall lipids such as di-acyltrehaloses (DATs) and poly-acyltrehaloses (PATs) essential for virulence (19–23), for regulation of the enduring hypoxic response and respiration, secretion of virulence factors (24) and most importantly for arresting the maturation of phagosomes, an event that is crucial for the intracellular survival of mycobacteria (25). In fact, a single point mutation in the *phoP* gene contributes significantly to the attenuation of *M. tb* H37Ra, the isogenic counterpart of the virulent *M. tb* H37Rv laboratory strain (23, 26, 27). Studies have demonstrated improved clearing of H37Ra as compared to H37Rv in infected alveolar macrophages and murine models, possibly due to the inability of H37Ra to arrest the movement of lysosomes in the outer periphery of the cell and towards the nucleus, resulting in mycobacterial cell death (24, 26). Further, Abramovitch et al., have identified an acid and phagosome regulator locus that is activated by PhoPR under acidic pH environment to secrete molecules/peptides that inhibit phagosome maturation and prevent killing of the pathogen (17). Notably, an *M. tb phoPR* knockout strain is attenuated for intracellular growth in human and mouse models of infection (22, 23), and is a component of MTBVAC-the first live attenuated *M. tb* based vaccine currently in clinical trials (28).

The traditional belief that TCS signaling pathways are linear has now been replaced with concrete evidence from several studies showing extensive cross-talk between cognate and non-cognate pairs of SKs and RRs (29, 30). Furthermore, the emerging network is no longer restrictive to TCS components but extends to *M. tb* STPKs as well (8, 31, 32). Conceivably, these added layers of regulation allow fine tuning of regulatory pathways critical for pathogenesis and intracellular survival. For example, PhoP RR interacts with DevR RR of the DevRS/T signaling system implicated in regulating mycobacterial dormancy response (16). The interplay of both PhoP and DevR DNA-binding regulatory proteins is believed to regulate its transition into a state of dormancy or persistence. While there is a considerable amount of information on the regulatory effects of PhoP RR, very little is known about the PhoR sensor kinase, the network of its cross-interacting partners and the underlying mechanism of activation.

In this study, we have studied the potential of PhoR as a unique target for developing novel anti-tubercular drugs. We demonstrate that in addition to its cognate RR, PhoR sensor kinase directly phosphorylates DevR, enabling the flow of information between the PhoPR and DevRS/T regulatory pathways. Furthermore, PhoR itself undergoes Ser/Thr phosphorylation by *M. tb* PknK, a growth regulatory STPK. We rationalized that inhibition of PhoR autophosphorylation could result in significant perturbation of downstream signaling pathways, and aid in the identification of drugs that could act on active as well as dormant bacilli. Towards this end, we developed a high throughput assay to screen for small molecule inhibitors of PhoR autophosphorylation. Of the several hits obtained, we confirmed and validated the inhibitory effect of Tamoxifen (TAM) on mycobacterial growth and PhoPR function and discussed its potential applicability as an anti-TB drug.

## Results

### Non-cognate interactions of *M. tb* PhoR sensor kinase with TCS and STPK proteins

Given the role of PhoPR TCSs in mycobacterial virulence (20, 27, 33–35), it is pertinent to understand the interactions of PhoP and PhoR proteins with other signaling proteins. Since PhoP interacts with DevR RR of the DevRS/T TCS system (16), we sought to investigate whether DevR RR is a part of the PhoR signaling cascade. For this, we purified recombinant His-tagged PhoR SK (cytosolic domain; ∼27 kDa) and full-length DevR RR (∼25 kDa) proteins to homogeneity and performed *in vitro* kinase assays as a function of time. The functional activity of PhoR was confirmed using an ATP depletion assay as per the manufacturer’s instructions (Fig. S1A). Figure 1A depicts the autophosphorylation of PhoR and the subsequent phosphotransfer from PhoR∼P to DevR RR. Densitometric analysis of the autoradiogram revealed a progressive increase in DevR phosphorylation over 30 min confirming a direct role of PhoR in activating DevR RR (Fig. 1B). Equal loading of proteins in all lanes was confirmed through CBB staining (Fig. 1A, lower panel). Furthermore, the binding affinity of PhoR and DevR interaction was calculated through MST and was found to be 683 ± 104 nM (Fig. S2). Recent studies have revealed the involvement of *M. tb* PknK in regulating PrrA and MtrA RRs, which are essential for mycobacterial growth and survival (11, 12, 32). However, very little is known about the *O*-phosphorylation of HKs through the Ser/Thr protein kinases. Given that the PhoPR system is activated under acidic pH conditions and the fact that PknK regulates growth under acidic pH (36), we wanted to investigate possible interactions between STPK PknK and PhoR HK. Towards this, *in vitro* kinase assay was done with purified His-tagged recombinant PhoR and full-length PknK wild type and mutant proteins followed by visualization with ProQ™ Diamond phosphoprotein stain. The functional activity of PknK was confirmed using an ATP depletion assay (Fig. S1B). Because the ProQ stain is specific for Ser/Thr phosphorylation (37), no signal was observed on the PhoR protein when incubated alone with ATP for 60 min, a condition where it can undergo autophosphorylation at a conserved Histidine residue. (Fig. 1C, Lane 1). Interestingly, we observed robust phosphorylation of PhoR HK in the presence of wild-type PknK protein (Fig. 1C, Lanes 2-4). The signal intensity increased over a duration of 60 min, indicating stable phosphorylation of PhoR (Fig. 1D). To further confirm whether the signal is PknK specific, we performed the assay with the phosphorylation defective mutant of PknK (K55M mutant) previously shown to be kinase-dead (38, 39). As expected, no signal was observed on PhoR in the presence of the mutant kinase (Fig. 1C, Lane 5), thus confirming that PhoR is subjected to phosphorylation via PknK. Such multi-site (His/Ser/Thr) phosphorylation of HKs is not well characterized in mycobacteria and may have implications on the regulation of PhoR structure and function. These results place PhoR at the centerstage of the *M. tb* signaling landscape and highlight its essential role in mycobacterial pathobiology.

**Fig. 1.**
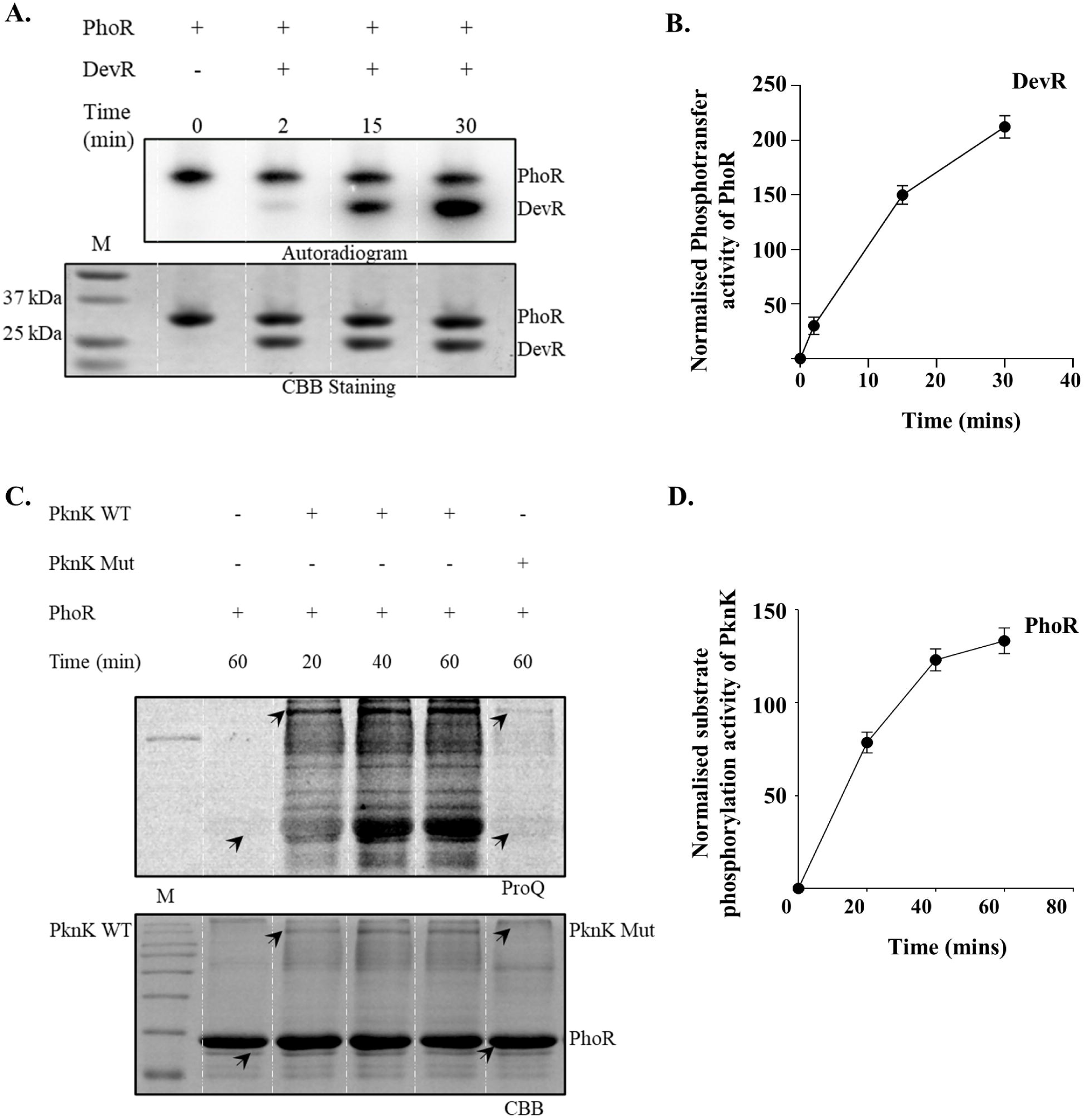
Non-cognate signaling interactions of *M. tb* PhoR sensor kinase. *In vitro* phosphotransfer assay of autophosphorylated sensor kinase PhoR with **(A)** DevR RR. PhoR was autophosphorylated for 1 h in the presence of γ-^32^P labeled ATP and phosphotransfer reactions to DevR RR were performed for different time points as indicated. **(B)** Densitometric analysis of the kinetics of DevR phosphorylation by PhoR using the Image J software. Mean ± SD from three independent replicates are plotted. *In vitro* kinase assays of PhoR with **(C)** STPK PknK. Both PknK and PhoR kinases were incubated together in the presence of ATP and phosphorylation was followed for different time points as indicated. Visualization was done using ProQ™ Diamond phosphoprotein stain. Top, Autoradiogram or ProQ stained gel; bottom, CBB stained image of the same gel. M, protein marker. **(D)** Densitometric analysis of the kinetics of PhoR phosphorylation by PknK using the Image J software. Mean ± SD from three independent replicates is presented.

### Development of a high-throughput screening assay for PhoR autophosphorylation inhibitors

The interconnectedness of the PhoR sensor kinase with cognate and non-cognate TCS systems, and the STPK signaling proteins provide the impetus to investigate PhoR as a potential target for the development of anti-tubercular drugs. For this, we developed a high-throughput 96-well plate assay to screen for compounds that inhibit PhoR autophosphorylation. We optimized a high throughput screening (HTS) platform specifically targeting the autokinase activity of PhoR. The schematic workflow of the HTS assay is shown in Fig. 2A. Autophosphorylation of PhoR-GFP was scored by measuring the intensity of spots on the autoradiogram (Fig. 2B and 2C). The Z-score of the radioactive dot blot assay developed was calculated to be 0.95, confirming no overlap between the positive and negative controls. In this experiment, we used GFP-tagged PhoR to normalize the protein amounts on the membrane, to account for well-to-well variation. Reactions in the presence of EDTA and DMSO served as negative and positive controls, respectively. The assay was validated multiple times and proved to be fast, sensitive, and reproducible.

**Fig. 2.**
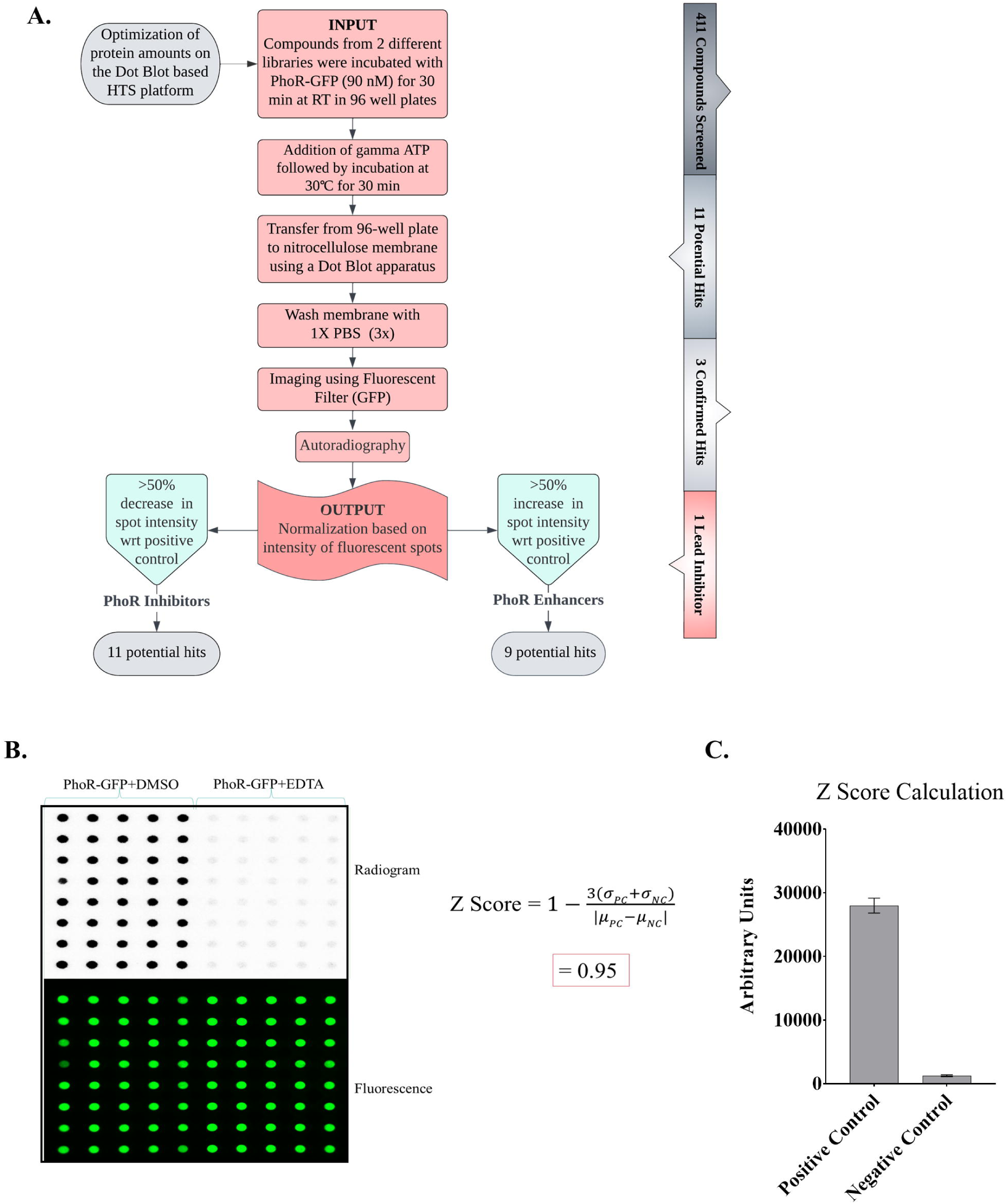
Development of high-throughput screening assay. **(A)** Schematic of the workflow for development of the HTS platform for PhoR-GFP autophosphorylation inhibitors. **(B)** represents autoradiogram (upper panel) after autophosphorylation of PhoR-GFP normalized with the corresponding fluorescence spots (lower panel). **(C)** represents the densitometric analysis done to calculate the Z-score. PhoR-GFP+EDTA and PhoR-GFP+DMSO served as the negative and positive controls, respectively.

### Screening of compounds and validation of hits

Two different chemical compound libraries (NIH-NCATS oncology set and Sigma LOPAC^1204^) were used, and ∼500 compounds were screened for probable inhibitors of PhoR autophosphorylation (Fig. 3A and 3B). Screening was done in duplicates and the readout was obtained by measuring spot intensities as described in Methods. Compounds inhibiting more than 50 % of autophosphorylation *in vitro* were classified as inhibitors. We identified 11 potential hits, which were taken forward for validation (Table 1). Colored compounds with absorption spectra coinciding with that of GFP, for example, Doxorubicin (with an Absorption maximum of 496 nm) were classified as false positives as they yielded very high fluorescence values. Some compounds acted as chelators and hence were unfavorable for downstream assays, while many were not commercially available for further testing. Only 3 compounds displayed more than 50 % inhibition in the dot-blot assay without interfering with the assay. Coincidentally, 2 of the 3 compounds originating from two different libraries were the same, a well-known breast cancer therapeutic called Tamoxifen (TAM) (40) (Fig. 3C, *left panel*). The other compound was identified to be Actinomycin D (Fig. 3C, *right panel*). We confirmed the inhibitory action of TAM on PhoR kinase activity. As shown in Fig. 3D, autophosphorylation of PhoR was inhibited in the presence of TAM. Densitometric analysis revealed a ∼56 % reduction in PhoR autophosphorylation in the presence of TAM (Fig, 3D, Lanes 4 and 5) compared to PhoR alone (Fig 3D, Lane 1). Since Actinomycin D is a known carcinogen, further experiments were done with TAM only.

**Fig. 3.**
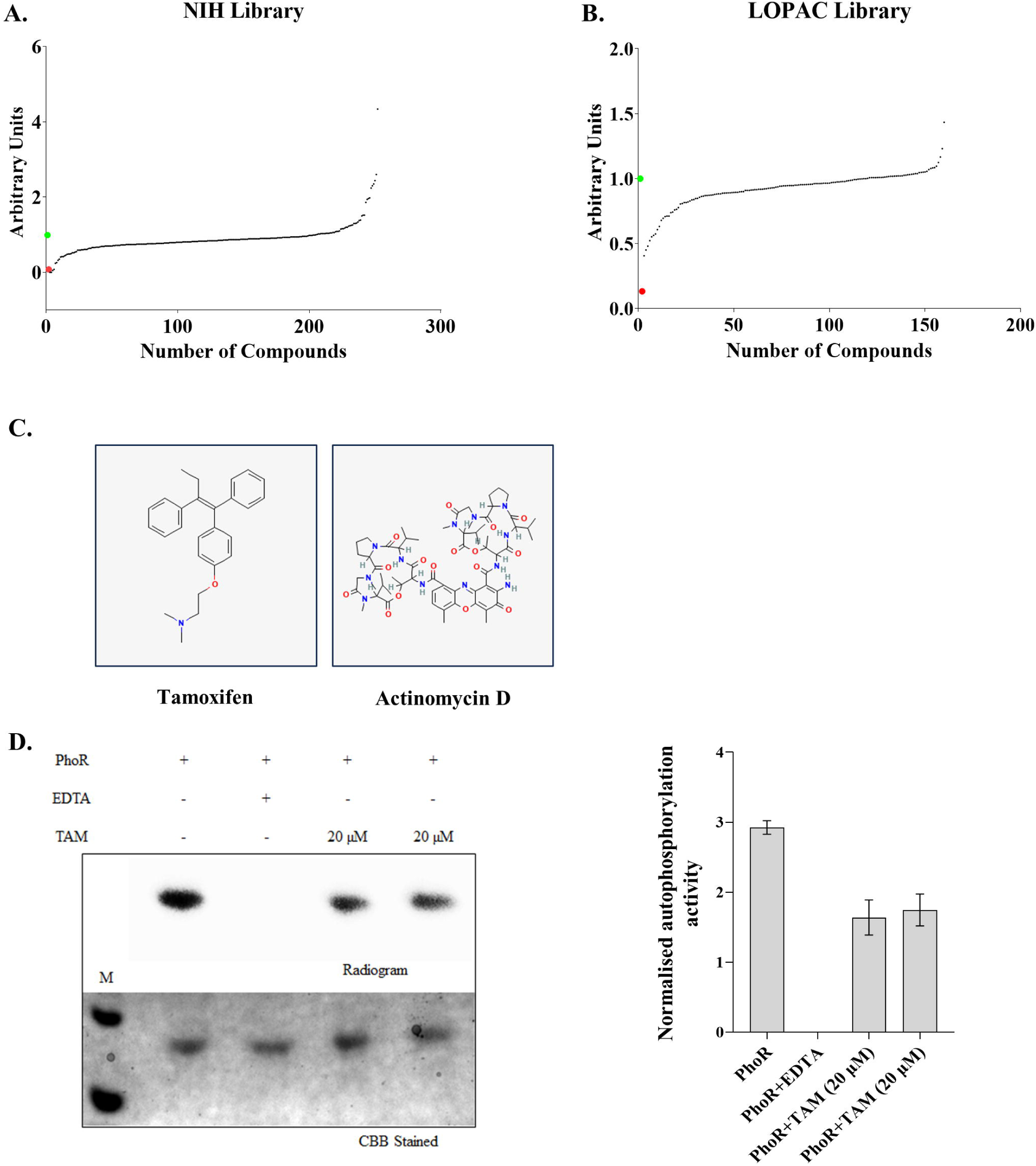
Screening and identification of compounds. Plot of autophosphorylation signal intensity normalized with the amount of protein against the compound number sorted in ascending order. **(A)** shows data of the compounds from the NIH Library (n=230 compounds) and **(B)** from the LOPAC library (n=181 compounds). Red and Green dot denote no autophosphorylation in the presence of EDTA (negative control) and autophosphorylation in the absence of any inhibitor (positive control), respectively. **(C)** Structure of Tamoxifen (TAM) *(left panel)* (PubChem CID 2733526) and Actinomycin D *(right panel)* (PubChem CID 2019). (D) Kinase assay of PhoR with TAM. Reaction in the presence of EDTA served as negative control (Lane 3). Assay was put up in duplicate with PhoR and 20 µM TAM, incubated for 60 min, and visualized by autoradiography (Lanes 4 and 5). Densitometric analysis of the autoradiogram is shown, highlighting the inhibitory action of tamoxifen.

**Table 1.** List of potential hits obtained in the HTS Assay.

| NSC<sup>a</sup> | % Inhibition<sup>b</sup> | Compound Name (CAS No.)<sup>c</sup> |
| --- | --- | --- |
| 180973 | 69 | Tamoxifen citrate (54965-24-1) |
| 73495 | 92 | B 132742 |
| 76455 | 58 | Peliomycin (1404-20-2) |
| 145366 | 96 | 2-Propanol, 1,1'-[(1-methylethylidene) bis(4,1-phenyleneoxy)] bis[3-[(1,1,3,3-tetramethylbutyl)amino]-, dihydrochloride |
| 173046 | 52 | Diphenyllead diacetate (6928-68-3) |
| 3053 | 100 | Actinomycin D (50-76-0) |
| 122819 | 100 | Teniposide (29767-20-2) |
| 123127 | 74 | Doxorubicin (25316-40-9) |
| 180973 | 77 | Tamoxifen citrate (54965-24-1) |
| --- | 60 | Protoporphyrin IX Disodium (50865-01-5) |
| --- | 55 | Caroverine (23465-76-1) |
| --- | 52 | Yohimbine (146-48-5) |

<sup>a</sup>Cancer Chemotherapy National Service Centre number is a compound identifier assigned by DTP to identify an agent or product.
<sup>b</sup> Percent inhibition calculated wrt positive control after normalization of protein amounts.
<sup>c</sup> Identity of compounds and CAS number.

### Binding studies of Tamoxifen to PhoR

The PhoR sensor kinase encoded by *Rv0758* gene, is a 485 amino acid long transmembrane protein that exists as a homodimer (41). Biochemical studies have established the role of a conserved histidine residue at position 259 in PhoR autophosphorylation and phosphotransfer of the signal (41). Understanding the binding between Tamoxifen and PhoR is important for understanding its specificity and efficacy and thus, we utilized Microscale Thermophoresis (MST) to determine the binding affinity of Tamoxifen with PhoR. We observed that tamoxifen binds to PhoR *in vitro* with a binding affinity (K_d_) of 108.5 ± 44 nM (Fig. 4A).

**Fig. 4.**
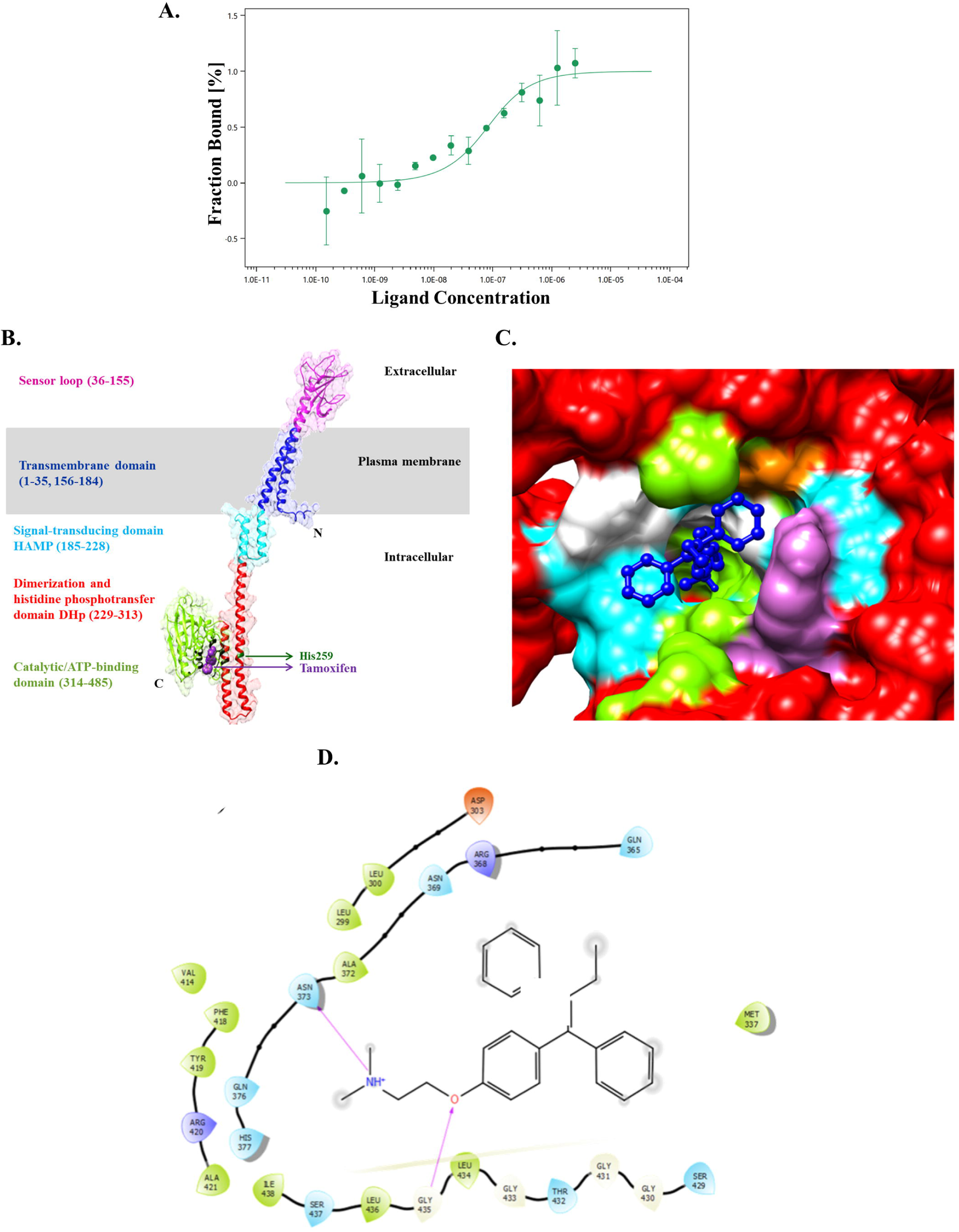
Binding interactions of Tamoxifen to PhoR. **(A)** Microscale Thermophoresis showing K_d_-Fit analysis of PhoR-GFP with Tamoxifen. The experiment was performed in triplicates and one representative graph is presented. **(B)** PhoR predicted structure model and binding site. Ribbon diagram representation of PhoR model structure obtained from AlphaFold database highlighting the different domains along with amino acid residue numbers, His259 and Tamoxifen bound at Site1. Surface representation of PhoR (red) – Tamoxifen (dark blue ball and stick model) docked complex **(C)** highlighting interacting residues (hydrophobic-green, polar-cyan, positively charged-purple, negatively charged-orange and glycine-white) and **(D)** Protein-ligand interactions within 0.5 nm are shown (Hydrogen bonds-pink lines).

To gain further insight into the structural aspects of ligand binding to PhoR, we used computational tools. Three-dimensional structure of PhoR was obtained from AlphaFold database. Structure validation statistics revealed that the 485 amino acid predicted structure of PhoR was of very good quality having ERRAT quality factor of 94.27% and 96.7% residues in the favorable region of the Ramachandran plot. Since binding of PhoR to Tamoxifen compromises its autophosphorylation activity, it was important to understand the associated structural changes. PhoR domain architecture along with surface representation of PhoR-TAM complex and interacting residues is shown in Fig.4B, 4C and 4D. Binding site prediction for kinase domain of PhoR spanning amino acid residues 256 to 470, resulted in three probable ligand binding sites i.e. Site 1 (residues 296-462), Site 2 (residues 242-445), and Site 3 (residues 260-431). Molecular docking of PhoR with TAM at all three sites revealed Site 1 having the most negative binding free energy value (ΔG) of ∼ -52.68 kcal/mol as opposed to ∼ -32.55 kcal/mol at Site 2, and ∼ -30.91 kcal/mol at Site 3. Our analysis suggests that TAM prefers to bind unphosphorylated PhoR at Site 1 (-52.68 kcal/mol). We noted that in comparison to unphosphorylated PhoR, PhoR∼P showed a significant reduction in free energy value (-1.25 kcal/mol). These observations corroborate our experimental data, wherein we show that TAM inhibits PhoR autophosphorylation.

### Sensitivity and Specificity of Tamoxifen

The IC50 value of Tamoxifen for PhoR was determined using a dot-blot assay investigating the effect of varying concentrations of TAM on the inhibition of PhoR autophosphorylation. As shown in Fig 5A and 5B, 10 µM Tamoxifen reduced the autokinase activity of PhoR by ∼40 % and thus IC50 for TAM was estimated to be in the range of 10-100 µM.

**Fig 5.**
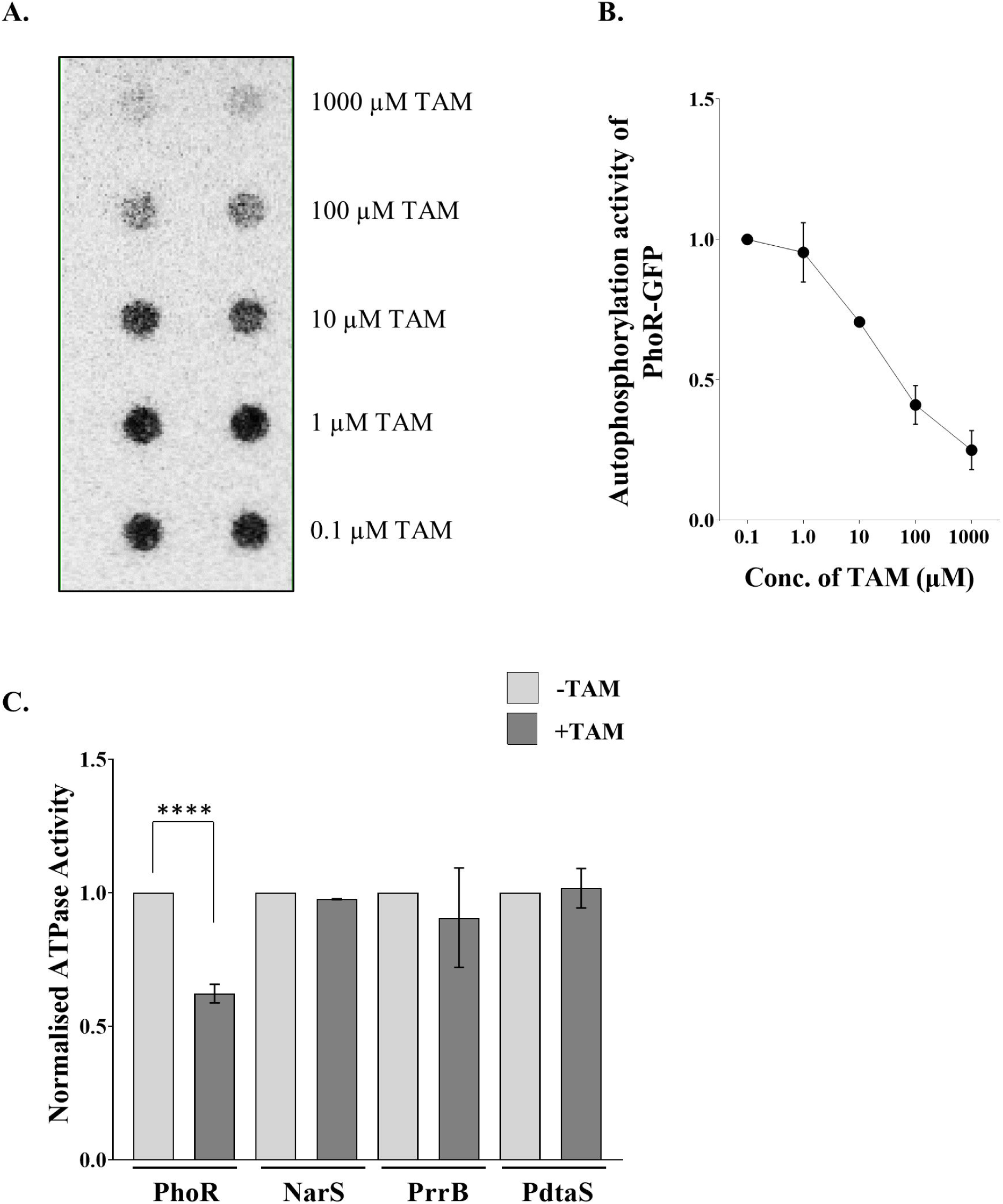
Sensitivity and specificity of Tamoxifen. **(A)** Dot Blot with increasing concentration of Tamoxifen performed in duplicates for IC50 estimation. **(B)** densitometric analysis of the autoradiogram presented in Fig. 5A. **(C)** Effect of TAM on autophosphorylation activity of a subset of *M. tb* HKs. ATP depletion assays and normalized ATPase activities from radioactive kinase assays are plotted for the specified HKs in the presence and absence of 10 µM Tamoxifen. Data for every HK is presented w.r.t minus drug control (set at 1.0).

To address the question regarding the specificity of TAM for PhoR HK, we determined the effect of TAM on three other HKs, namely PrrB, NarS and PdtaS using an ATP determination assay as per manufacturer’s protocol. While TAM exerted inhibitory effects on PhoR ATPase activity, there was no significant inhibition observed with other HKs tested in the assay (Fig. 5C). These data are supported by independent autoradiography experiments, wherein the signal intensity obtained with HKs in absence or presence of TAM was measured by densitometry and plotted (Fig. S3). Similar results were obtained for PdtaS and SenX3 with radioactive kinase assays. Thus, among the five HKs tested in this study, TAM directed autoinhibiton of PhoR only. Predictably, it follows that in the absence of PhoR autophosphorylation, there would be reduced downstream PhoR-mediated phosphotransfer and activation of PhoP, thus impacting the expression of PhoPR regulon genes.

### Tamoxifen inhibits PhoR-mediated signaling *in vivo*

Since most of the major frontline drugs for different strains of TB (like Isoniazid and Rifampicin) have MIC values of 5-10 µg/ml (42) to inhibit or slow down the growth of drug resistant strain of mycobacteria, we used 10 µg/ml of Tamoxifen to check its effect on the growth of *M. bovis* BCG. Earlier, Jang et al., have reported the MIC for TAM in the range of 6.75 µg/ml to 12 µg/ml (43). Activation of PhoR is facilitated by low pH (17, 44, 45), therefore we conducted the growth experiments at acidic pH (pH 5.5) and neutral pH (pH 7.0). As expected, the trends were similar for the control and vehicle (DMSO) treated cells at both pH 7.0 (Fig. 6A) and pH 5.5 (Fig. 6B), with the overall growth being less under conditions of acidic pH. In comparison to the vehicle control, growth was significantly inhibited in the presence of tamoxifen at pH 5.5 with ∼50 % reduction in the optical density (Fig. 6B).

**Fig. 6.**
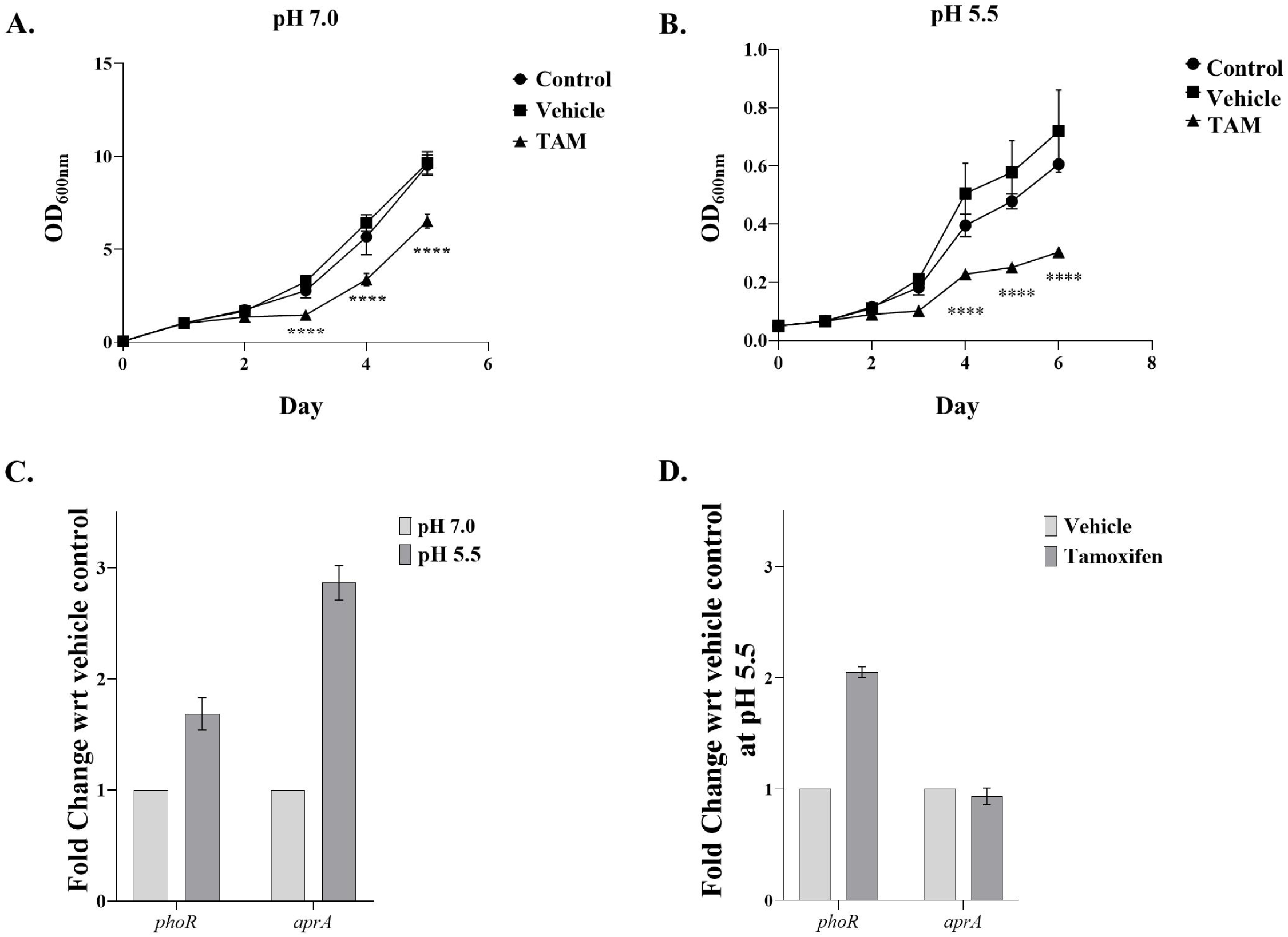
Effect of Tamoxifen on *M. bovis* BCG growth and PhoPR downstream signaling. Plot of optical density at 600 nm of *M. bovis* BCG versus time (days) at (**A)** pH 7.0 and **(B)** pH 5.5. Solid circles and squares represent growth in the absence (control) and presence of DMSO (vehicle), respectively. Solid triangles represent mycobacterial growth in the presence of Tamoxifen (TAM). The difference between the growth of control cultures versus *M. bovis* BCG exposed to the inhibitor is statistically significant (n=3, \*\*\*\**P* < 0.001). **(C)** and **(D)** represents real time PCR analysis of *phoR* and its downstream target gene *aprA* at pH 5.5 and pH 7.0. Fold change is reported w.r.t vehicle control (baseline as 1.0). Mean ± SD from 3 independent experiments is plotted.

**Fig. 7.**
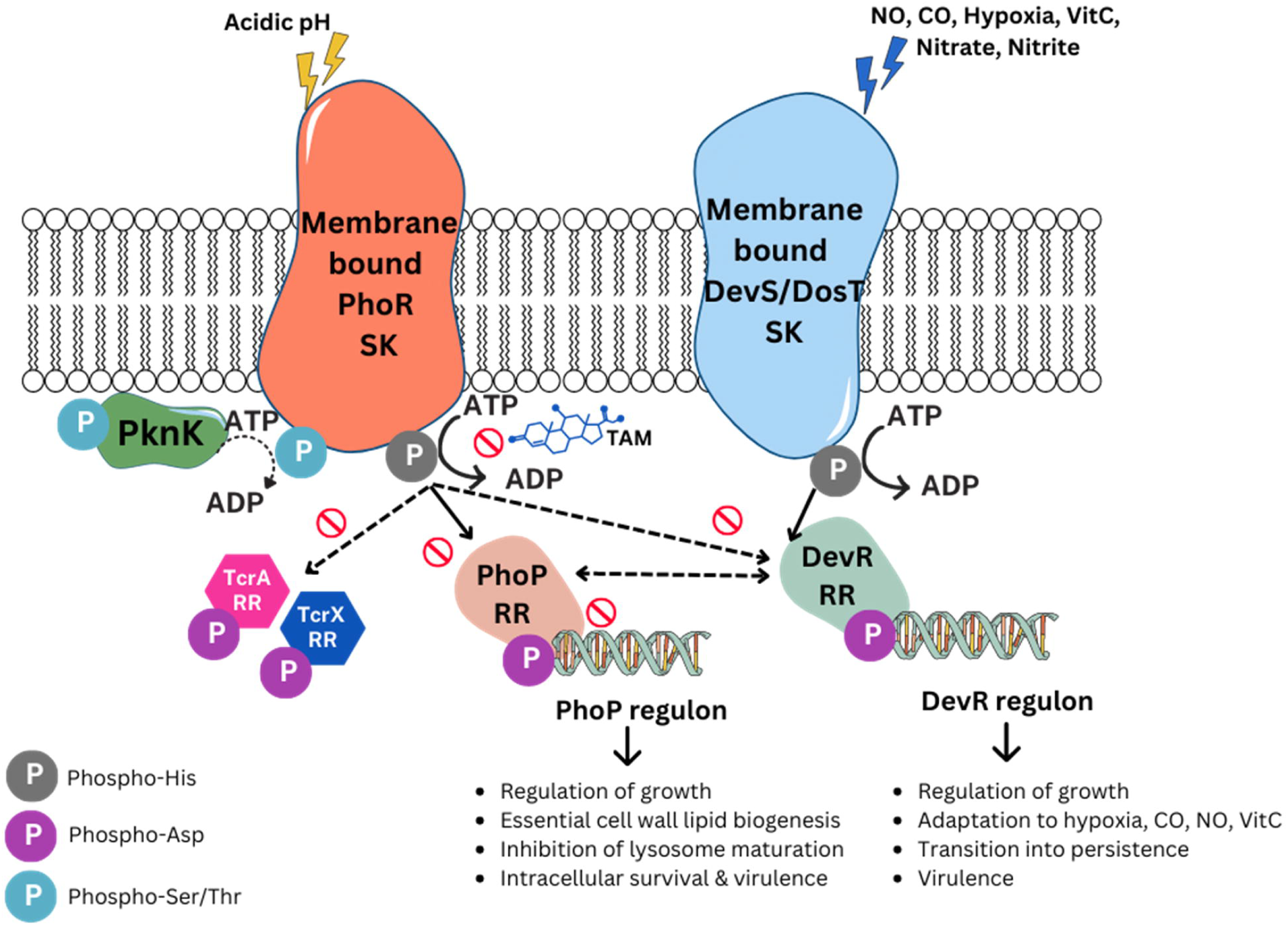
A proposed model for Tamoxifen action on *M. tb* PhoPR TCS system. A model is proposed based on the data obtained from this study and previously published literature (43, 50). A comprehensive view of TCS crosstalk was reported previously (8, 30). Here, we have depicted cognate (solid arrows) and non-cognate (dashed arrows) interactions of the PhoPR TCS system only. PhoR kinase senses external stimuli such as acidic pH (17, 44, 45) and undergoes autophosphorylation at His259 residue (41). The signal is then transferred to PhoP through a phosphotransfer reaction, thereby activating PhoP, which binds to the promoter regions of PhoP regulon genes. Tamoxifen binds to PhoR and blocks PhoR autophosphorylation and, thus, inhibits downstream activation of PhoP, resulting in abrogation of the upregulation of *aprA*, an acid-induced PhoP target gene (17). Since PhoP and DevR interact and coregulate DevR regulon genes (16), the inhibitory effect of tamoxifen on the dormancy response is plausible, although other independent pathways feeding into DevR must also be considered. PhoR is also subjected to *O*-phosphorylation via PknK, a soluble/membrane-associated STPK; however, the downstream effects are yet unknown. It should be noted that the effect of tamoxifen on other TCS pathways cannot be excluded and necessitates further investigations.

RNA isolated from *M. bovis* BCG cultures on day 6 of treatment was analyzed for the expression of PhoPR-regulated genes. It is known that PhoP∼P binds to *aprA* promoter causing upregulation of *aprA* expression at pH 5.5 (17). The rationale was that if Tamoxifen inhibited PhoR activation, the downstream signaling would be affected and hence, *aprA* induction would be abrogated. Quantitative gene expression analysis revealed induction of *phoR* (∼1.8-fold) and *aprA* transcripts (∼2.9-fold) at pH 5.5 when compared to pH 7.0 (Fig. 6C) confirming the activation of the PhoPR TCS and downstream regulation at acidic pH in the absence of the drug. Interestingly, we noted an increase in *phoR* expression at pH 5.5 in the presence of Tamoxifen as well (Fig. 6D). This may be attributed to the autoregulatory effects of the PhoP response regulator already present in the cell (18). However, downstream target gene induction requires PhoP phosphorylation and activation via PhoR (18). Despite the increased number of *phoR* transcripts, no induction was observed for *aprA* indicating the absence of PhoPR activation. Taken together, these data suggest a plausible mechanism for Tamoxifen action in mycobacteria.

## Discussion

Two-component signaling proteins are believed to be desirable drug target candidates owing to their integral role in driving adaptive responses to extracellular or intracellular cues, which include antimicrobial resistance, virulence regulation, bacterial growth, and survival. To date, multiple TCS system inhibitors with diverse chemical and structural properties have been biochemically characterized (46). While both the sensor kinase and the DNA-binding response regulator are druggable targets, there is an inherent benefit of shutting a signaling pathway by inhibiting the histidine sensor kinase, as the drug blocking them does not have to be taken up by the bacterium and the cascade can be inhibited at the first step. Moreover, the conserved features of HK domains may facilitate development of inhibitors possessing broad-spectrum antimicrobial activity.

In this study, we have focused on evaluating the potential of *M. tuberculosis* PhoR HK as a novel antimycobacterial drug target primarily due to its role in virulence (20), regulation of essential cell wall lipid synthesis (22, 33), and its complex network with non-cognate signaling proteins (30). It is interesting to note that the PhoPR and DevRS/DosT TCS systems converge at multiple points. Previously, it was shown that PhoP and DevR RRs form heterodimers to coregulate the dormancy regulon in H37Rv (16). Here, we demonstrate the cross-talk between PhoR sensor kinase and DevR response regulator, adding yet another layer to the PhoPR-DevRS regulatory network. We also show STPK-mediated phosphorylation of a mycobacterial histidine kinase. PhoR is subjected to *O*-phosphorylation by *M. tb* Ser/Thr Protein kinase K (PknK), a soluble / membrane-associated kinase, that has multiple TCS RRs as targets (32). Although the effect of phosphorylating PhoR at multiple amino acid sites are not fully understood yet, the data presented here clearly creates value addition to the potential of PhoR SK as the single most connected HK involved in regulating mycobacterial pathogenesis and virulence. Our goal in this study was three-fold: (i) to develop and optimize a function-based high-throughput screening assay for PhoR autokinase activity; (ii) to screen for small molecule inhibitors that abolish PhoR autophosphorylation and; (iii) to validate the effect of inhibitor on downstream regulation of gene expression.

Histidine kinases being multifunctional enzymes can autophosphorylate, phosphotransfer, and dephosphorylate their cognate RR (or other substrate proteins) (46). All three enzymatic activities are possible points for targeting inhibition. Several studies have identified HK inhibitors and investigated their suitability for anti-TB therapy (reviewed in 46). Except for inhibitor molecules like HC102A, HC103A, and BTP15, which inhibit the autokinase activity of DevS/DosT (47), and MprB sensor kinases (48), respectively, the direct inhibition of PhoR function has not been investigated. In a study, Ethoxzolamide (ETZ) has been proposed to indirectly inhibit the PhoR sensing mechanism by targeting cell surface carbonic anhydrases that may modulate the local extracellular environment (49); however, no experimental evidence supports this model. Recently, Watson et al., reported that ETZ lowered the expression of target gene *aprA* and reduced CFU in a cell line infection model (18); however, no mechanism for ETZ action was proposed.

In our initial experimental approach, we examined PhoR-DevR protein-protein interaction using Microscale Thermophoresis (MST) in a HTS setting. Our goal was to identify compounds that disrupt HK-RR interaction and, thus, can potentially inhibit the phosphotransfer reaction from PhoR to DevR. However, the Z score of the MST-based assay was poor (∼0.6), and even after several rounds of optimization, could not be improved (Fig. S4A). Although we obtained 14 potential inhibitors of PhoR-DevR interaction after screening 722 compounds (Fig. S4B), none of them inhibited the phosphotransfer reaction in an *in vitro* kinase assay (Fig. S4C). With our modified approach targeting autophosphorylation inhibition, we were able to identify PhoR autokinase inhibitors as well as molecules that enhanced PhoR autophosphorylation. In principle, enhancing SK activity could also be an effective mechanism to dysregulate TCS pathways, trapping the kinase in an “On” conformation; however, in this study, we have focused on PhoR autokinase inhibitors only. Subsequent analysis and after application of cut-off value filters, only 2 hits were obtained. Identification of the same compound, Tamoxifen, from 2 different compound libraries; plate 4888 of the mechanistic set and plate 4891 of the oncology set of the NIH library raises the confidence of our data.

Tamoxifen is a clinically used anti-cancer drug that has been recently studied for its applicability in fighting antibiotic-resistant TB infections. Boland et al., have shown that tamoxifen is effective against clinical isolates of Multi-Drug Resistant (MDR) TB and inhibits bacterial growth in primary human macrophages; however, the underlying mechanism of tamoxifen action remains unknown (50). Although the direct antibacterial effects of tamoxifen against intracellular pathogens have been documented earlier (31), its potential to act as a host-directed therapeutic (HDT) for TB is just beginning to be investigated (40). HDT approaches have recently sparked interest as assistive adjunct therapies for TB treatment. The best-known target for tamoxifen is the host ER (40). Additionally, tamoxifen modulates other host pathways, such as autophagy and lysosome function (43, 50). Published data supports the notion that an increase in lysosomal activation by tamoxifen equips the host to control intracellular replication of pathogens such as *M. tb* (43, 50).

To the best of our knowledge, no mycobacterial target for tamoxifen has yet been reported. Given the fact that PhoPR TCS is responsible for sensing acidic pH within the macrophages and is necessary for intracellular survival, we propose that tamoxifen inhibits activation of PhoPR TCS by specifically inhibiting the autokinase activity of PhoR sensor kinase, thereby disrupting the downstream PhoPR signaling leading to decreased survival of the bacteria in acidic conditions (Fig. 6). The significant inhibition of growth observed in the *M. bovis* BCG cultures exposed to tamoxifen along with the concomitant inhibition of *aprA* expression, a PhoP-regulated target gene implicated in adaptation of mycobacteria to the acidic environment found in the phagosomes lend strong support to our model. The arrest of phagosome maturation is a key survival strategy of *M. tb,* that is mediated by the PhoPR TCS system. Thus, it is very likely that the effects exerted by tamoxifen on PhoPR signaling would lead to the host having an upper hand causing lysosomal activation and intracellular death of mycobacteria.

As mycobacteria possess multiple TCS systems and that many exhibit multiplicity in their interactions, inhibitors of the highly conserved active sites of HKs are expected to cause perturbations in multiple signaling pathways compromising the ability of the bacteria to rapidly adapt to environmental changes including those encountered in the host during infection. This may hold true for the PhoPR TCS considering its network with distinctly different signaling pathways. Arguably, in an intracellular environment primed with such promiscuous interactions, TCS inhibition may not be bactericidal, but it is highly likely to reduce pathogenic fitness. Overall, the data presented in this study highlights the value of PhoR as a novel drug target and shows how inhibition of PhoR via tamoxifen could prove to be a useful step towards understanding and devising better drug regimen to cure and eradicate TB.

## Experimental Procedures

### Strains, plasmids, and culture

The strains and plasmids used in this study are listed in Table S1. Luria Bertani (LB) broth and/or LB Agar was used to grow all the strains of *Escherichia coli (E. coli)* at 37 ℃. For protein purification, 2xYT was used to cultivate *E. coli* BL21 (DE3) cells. *Mycobacterium bovis* BCG cultures were grown in Middlebrook 7H9 liquid and 7H11 solid media supplemented with the OADC supplement (Middlebrook) at 37 ℃.

### Protein expression and purification

All proteins were purified from *E. coli* BL21 (DE3). Briefly, a single colony was inoculated in 5 ml of LB broth and cultured overnight at 37 ℃ with shaking at 180 rpm. 400 ml of 2xYT broth was then inoculated with 4 ml of overnight grown culture and grown at 37 ℃ with shaking till the OD_600_ reached 0.4 – 0.6. The culture was kept at 4 ℃ for an hour to achieve uniformity and induced with 1.0 mM Isopropyl-β-D-thiogalactopyranoside (IPTG) at 16 ℃ for 16 – 18 hrs. Subsequently, cells were harvested by centrifugation at 3500 rpm for 10 mins at 4 ℃. The cell pellet was resuspended in Lysis Buffer (50 mM Tris pH 8.0, 300 mM NaCl, 10 mM Imidazole and Protease Inhibitor cocktail) and lysed by sonication (20 mins, pulses of 2 sec ON and 2 sec OFF, 35 % amplitude). The soluble fraction was separated by centrifugation of the lysate at 13000 rpm for 30 min at 4 ℃. Supernatant was filtered using 0.2 µM filter and applied to His-Trap FF column pre-equilibrated with lysis buffer. The protein(s) were purified using Ni-NTA affinity chromatography. For elution of the protein, an elution buffer (50 mM Tris pH 8.0, 300 mM NaCl, 250 mM Imidazole) was used and fractions of 1 ml were collected. SDS-PAGE analysis was done to assess the purity of the protein. Pure fractions were then pooled and dialyzed in a dialysis buffer (50 mM Tris pH 8.0, 250 mM NaCl, 50 % glycerol) and stored at -20 ℃.

### *In vitro* kinase assay

To check for the kinase activity of PhoR SK, 90-100 pmoles of PhoR (or other HKs) was added in 20 µl reaction with 1x kinase Buffer (50 mM Tris pH 8.0, 20 mM MgCl_2_ and 1.0 mM DTT) with 50 µCi (ɣ-^32^P) labeled ATP for autophosphorylation for 1 h at 30 ℃. 10 – 12 µg of DevR was then added to the reaction mix as a substrate for 0 – 1 h at 30 ℃. The reactions were stopped by the addition of 5x SDS loading dye. The samples were resolved on 12% SDS-PAGE gels and analyzed using autoradiography. For kinase assays with PknK, both kinases were incubated in the presence of 1 mM ATP in 1x kinase buffer (25 mM HEPES pH 7.4, 15 mM MgCl_2_, 5 mM MnCl_2_, 1.0 mM DTT) and incubated for specified times. The reactions were stopped by adding 5x loading dye as before and were resolved on a 12 % SDS-PAGE gel. Ser/Thr phosphorylation was visualized using the ProQ™ Diamond Phosphoprotein Gel Stain (Thermo Scientific™) as per manufacturer’s instructions.

### High throughput assay to screen for small molecule inhibitors

To screen for compounds inhibiting autophosphorylation of PhoR, a 96-well plate assay using 96-well Dot Blot (Bio-Dot™, Bio-Rad Laboratories) was devised. 1 µg of PhoR-GFP was added to 1 ml of 1x kinase buffer and divided into 96-well containing 10 µM of different compounds each and incubated at RT for 30 mins. 10 µl 1x kinase buffer containing 1 µCi (ɣ-^32^P) labeled ATP was then added to each well containing PhoR-GFP with compounds and incubated at 30 °C for 1 h. Reaction mixes were then transferred onto a nitrocellulose membrane using the Dot-blot apparatus. The membrane was then washed with 1x PBS to remove unbound ATP and imaged for GFP fluorescence to check for equal loading of the proteins before analyzing for autophosphorylation using autoradiography.

### Z-score calculation

To calculate the Z-score of the assay, PhoR-GFP incubated with DMSO and and PhoR-GFP with EDTA were used as positive and negative controls, respectively. 48 wells of a 96-well plate were loaded with positive control (PC) and the remaining 48 wells with negative control (NC). 10 µl 1x kinase buffer containing 1 µCi (ɣ-^32^P) labeled ATP was then added to each well and incubated at 30℃ for 1 h. Reaction mixes were transferred onto the nitrocellulose membrane and analyzed as described above. Z-score was calculated using standard deviations and variances using the formula, 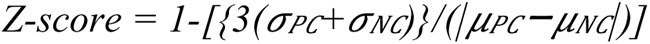.

### Microscale Thermophoresis

To determine the binding affinity of TAM and PhoR, TAM was procured (Selleck Chem, Cat No S1972), and dissolved in DMSO (10 mM). Serial dilutions of TAM were made in 16 different capillaries containing 50 µM PhoR-GFP (such that the fluorescence in the NanoTemper™ Monolith is in the range of 1200-1400 U). The capillaries containing the reaction mix were then subjected to MST analysis. PhoR-GFP and DMSO were used as controls. Data was analyzed using MO Affinity analysis software (NanoTemper Technologies).

### Protein structure retrieval and validation

Due to lack of availability of a crystal structure of PhoR, computationally predicted structure of PhoR was obtained from AlphaFold database (AF-P71815-F1) Structure was also validated using PROCHECK (52), which provides information about Ramachandran plot statistics, backbone conformation, stereochemical quality, and ERRAT program, which examines non-bonded atomic interactions in protein models (53).

### Molecular docking studies

Molecular docking was carried out for the cytosolic domain of PhoR with TAM. Prior to docking, the target protein was prepared using Protein Preparation Wizard (Schrodinger, LLC, New York, NY) which involved addition of the hydrogen atoms, assigning of bond orders, formation of disulfide bonds and removal of hetero-atoms followed by energy minimization and refinement. The ligand structure was prepared using the LigPrep module and binding site was determined using SiteMap module (54). Subsequently, molecular docking was done using extra-precision (XP) mode of Glide (55) and the binding free energy (ΔG bind) for each docked pose was further estimated using Prime MM-GBSA method (56). Docked complex structure of PhoR-Tamoxifen was used to further dock phosphoryl group at His259 residue.

### Phenotypic growth assay

*M. bovis* BCG cultures were grown in 10-20 ml of supplemented 7H9 media. Growth was followed in pH adjusted media (pH 5.5 and pH 7.0) from starting OD_600_ of 0.05 in the presence and absence of TAM (10 µg/ml) with DMSO as vehicle control. OD_600_ measurements were taken daily for 6 days. The cells were pelleted, washed with PBS, and stored at -70 ℃ for RNA isolation. Statistical analysis was done using one-way ANOVA (Prism 9.0 software).

### RNA isolation and Real-Time PCR analysis

For gene expression analysis, mycobacterial culture pellets were resuspended in TRIzol™ reagent (Thermo Scientific™) and processed for RNA isolation as per manufacturer’s protocol. Briefly, the suspension was lysed with 500 µl of sterile zirconium beads using a beat beater in 6 cycles of 30 sec at 30 Hz frequency. Beads were separated by centrifugation at 12000 rpm for 5 mins at 4 °C and supernatant was transferred in fresh 1.5 ml microcentrifuge tubes and further subjected to chloroform: isoamyl alcohol (24:1) extraction. RNA was precipitated from the upper aqueous phase with ice-cold ethanol. Supernatant was discarded and pellet was allowed to dry and dissolved in nuclease free water. The quantity of RNA was estimated using NanoDrop^TM^ and DNase treatment was done to get rid of any DNA contaminants. cDNA was synthesized using random hexamers supplied in the Verso cDNA synthesis kit (Thermo Scientific™).

Real-Time PCR analysis was done for selected genes using gene specific primers as described (18). Briefly, 100 ng cDNA prepared from vehicle control and TAM-grown cells, was taken and aliquoted into different wells of a 96-well plate for Real Time PCR. PCR Master mix (PowerUp™ SYBR™ Green Master Mix, Applied Biosystems) was added to each well along with the specific primer pair for each gene. The reactions were then subjected to 35 cycles of PCR and threshold cycle (C_t_) values were calculated for each gene. The calculated Ct value for each gene was normalized to *sigA* gene followed by that of the gene of interest in the vehicle control cells to determine fold change. Data from three independent biological replicates was analyzed and reported.

## Acknowledgments

We acknowledge the CIF facilities at the University of Delhi, South Campus and at the Department of Developmental Biology and Genetics, Indian Institute of Science, Bengaluru for infrastructure support. We also thank Devendra Pratap Singh, IISc for helping with the experiments and the members of TB MBSLab @SVC for critical review of the study.

## Author Contributions

AG designed the study, performed the experiments, analyzed data, and wrote the manuscript; MP performed the molecular docking studies and bioinformatic analysis; VM and DKS managed the project, supervised experiments, analyzed data and wrote the original manuscript.

## Funding and Additional Information

This work was supported by funds from the Department of Biotechnology (DBT), Government of India (DBT Grant No. BT/PR31937/MED/29/1404/2019) to V.M, and by BIRAC, DBT to D.K.S (BIRAC/PMU/2019/Sentinels-001) and the Department of Biotechnology (DBT), Government of India (DBT Grant No. BT/PR41993/MED/29/1559/2021) to DKS. A.G is grateful to DBT for JRF support.

