## supplementary_data for "Anti-cancer drug Tamoxifen interferes with *Mycobacterium tuberculosis* PhoPR mediated signaling and inhibits mycobacterial growth"

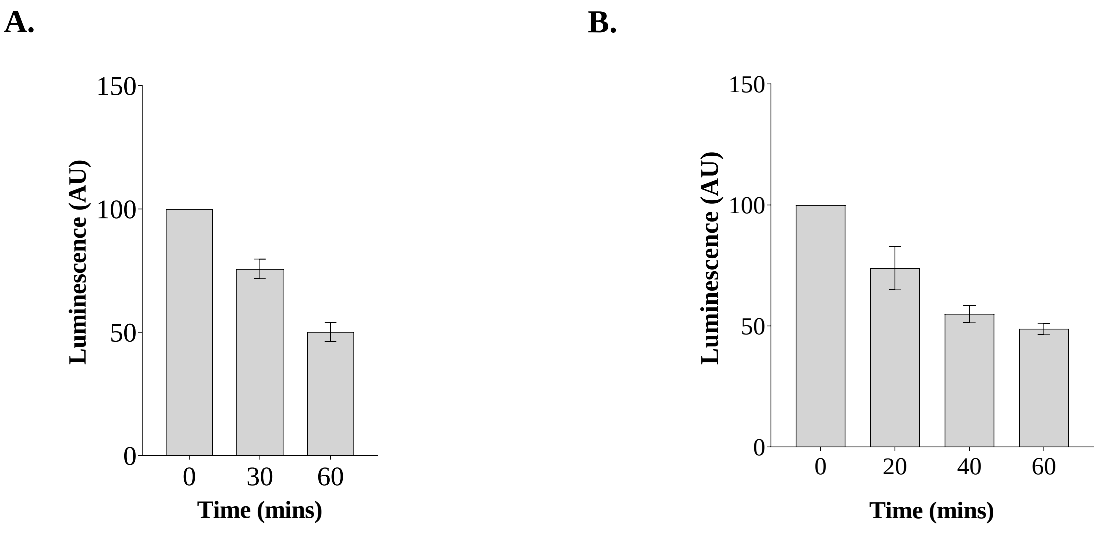


**Fig. S1. Functional activity of PknK and PhoR.**

ATP Depletion Assay was performed as per manufacturer's protocol to confirm that the purified proteins were active. (A) PhoR and (B) PknK were allowed to autophosphorylate for different time points as indicated and reaction mixture containing Luciferase and Luciferin was added. Luminescence was recorded based on the amount of ATP remaining after autophosphorylation by the respective kinase after mentioned time points (n =3).

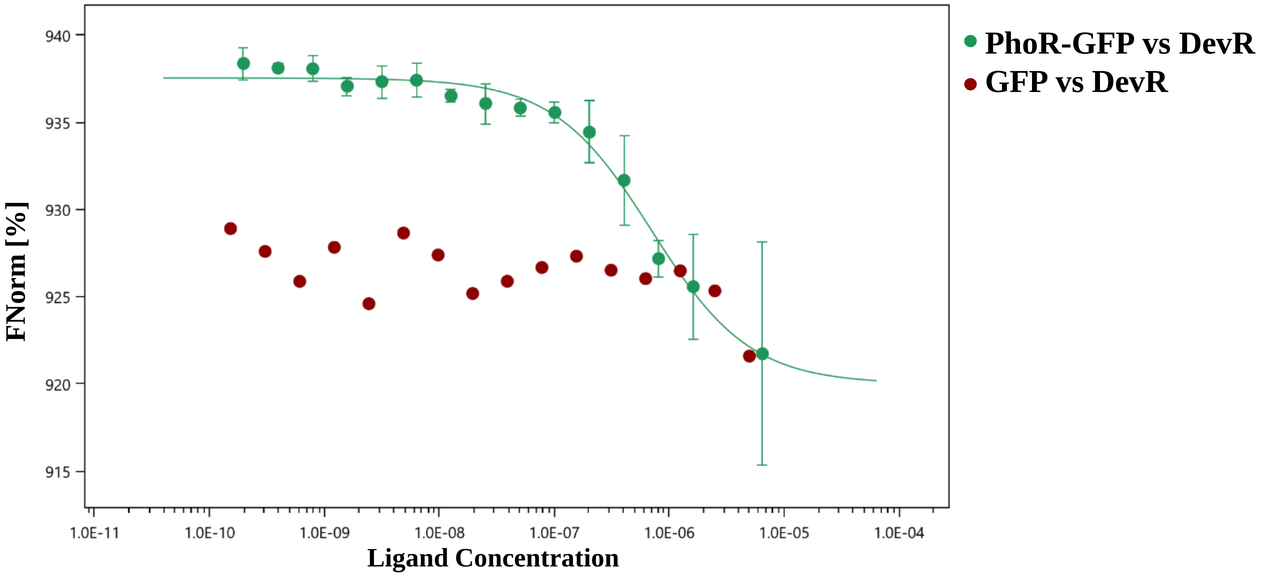


**Fig. S2. Binding analysis of PhoR-GFP and DevR.**

Microscale Thermophoresis showing $K_d$-fit analysis of PhoR-GFP with DevR. The experiment was done in triplicates and one representative graph is presented. Binding affinity of PhoR-GFP to DevR was calculated using MST as mentioned in the methods and was found to be 683± 104 nM.

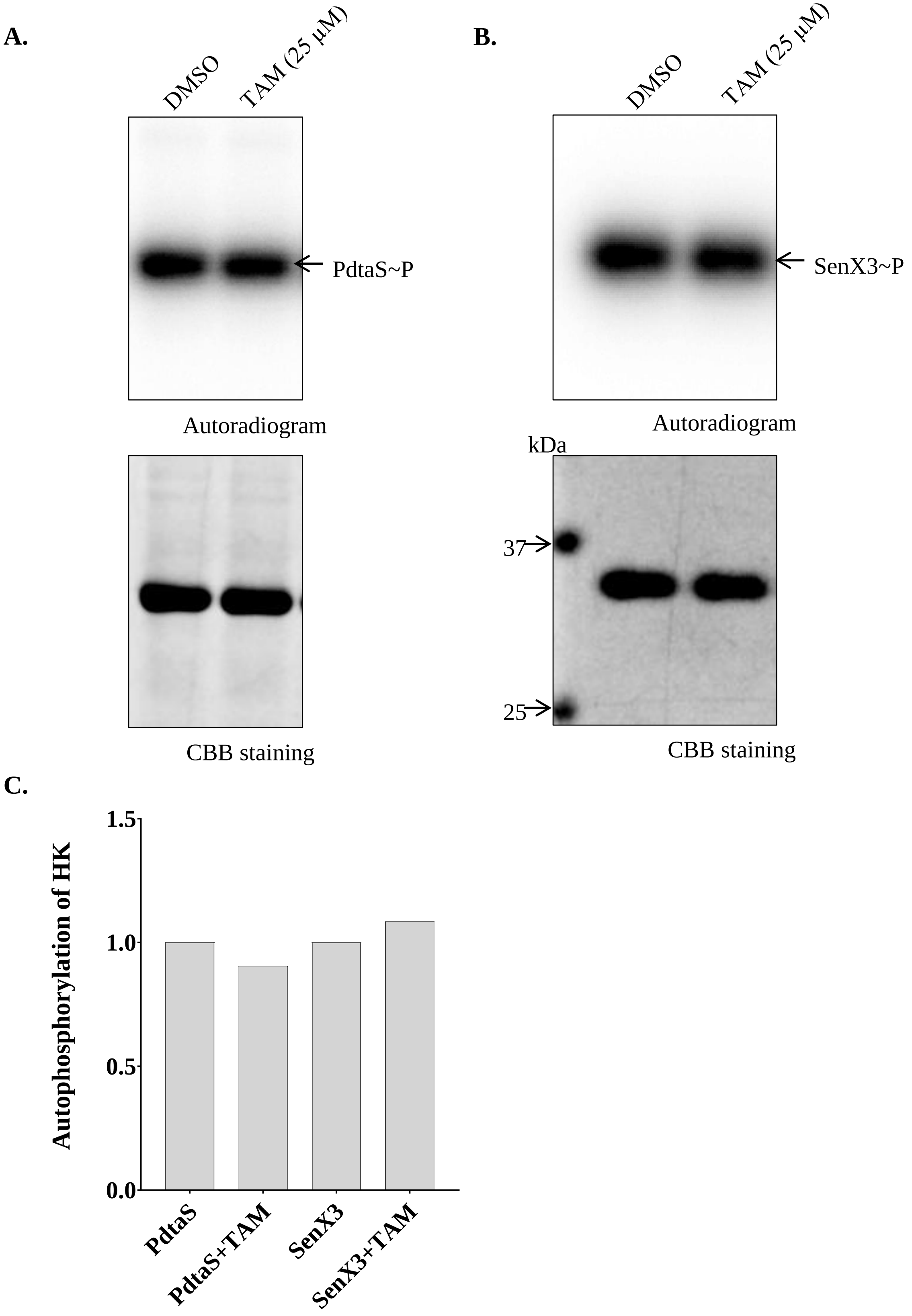


**Fig. S3. Autophosphorylation inhibition of HKs in presence of TAM.**

Radioactive *in vitro* kinase assays to check the autophosphorylation of (A) PdtaS and (B) SenX3 in the presence of 25 μM TAM and DMSO (vehicle control). (C) represents the densitometric plot of (A) and (B).

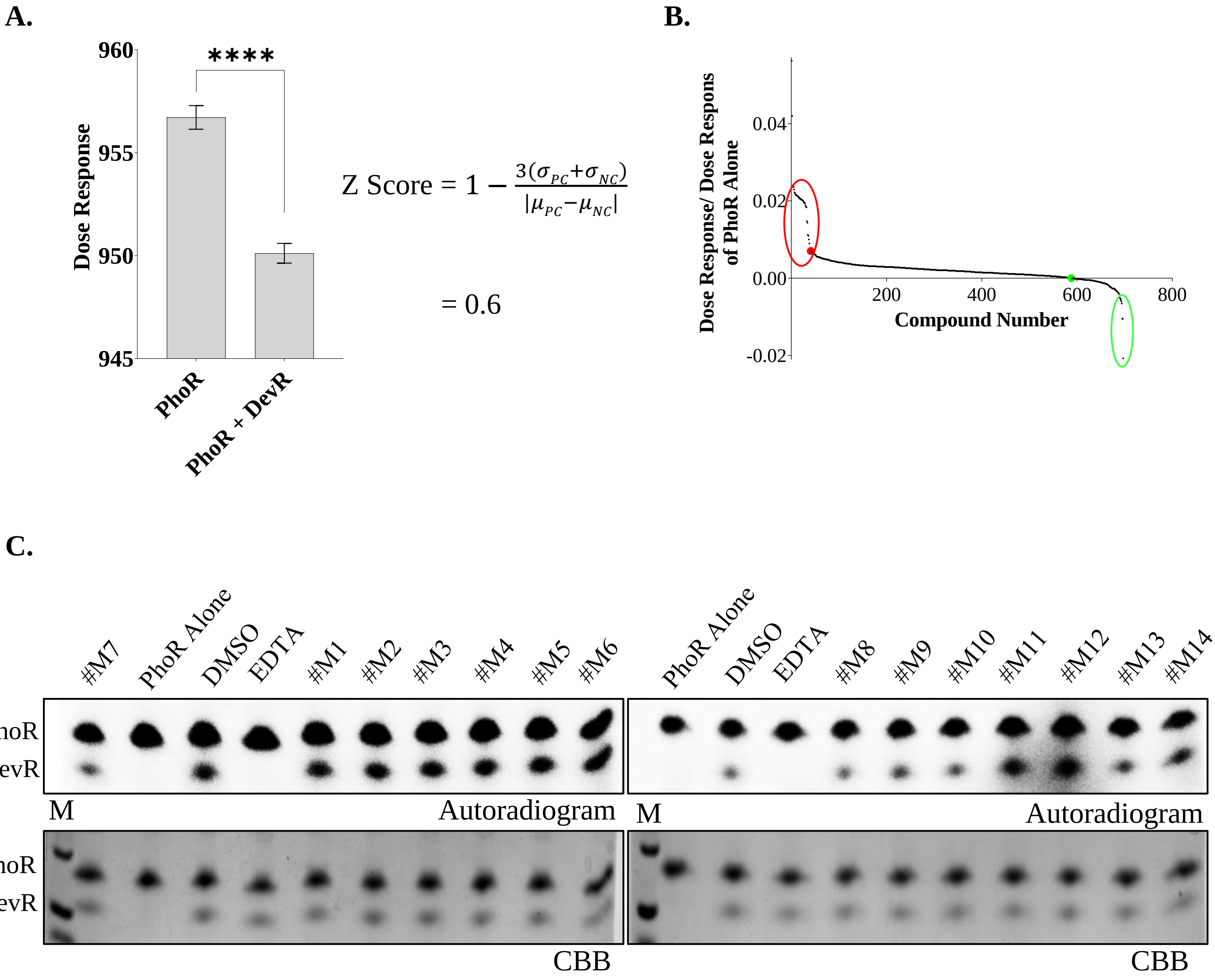


**Fig. S4. Z score calculation and results of HTS based on MST.**

(A) Z- score of HTS based on MST was calculated by recording the dose response of PhoR+DevR (Positive control) and PhoR alone (Negative Control) (n =16) (**** *P<0.001).* (B) Data from screening of 722 compounds. Red and green dots indicate negative and positive controls respectively. Red ring indicates inhibitors and green ring indicates activators/enhancers of PhoR~GFP – DevR interaction. (C) *In vitro* kinase assay to confirm the inhibition of phosphorylation of DevR phosphorylation by PhoR in the presence of different hits obtained. (#M1-#M14). EDTA served as positive control to show complete inhibition of phospho-transfer reaction.

**Table S1. List of Strains and Plasmids used in the study.**

| **Strain or plasmid construct** | **Description** | **Source or Reference** |
|---|---|---|
| *E. coli* DH5α | Δ*(argF-lac)169*, φ80d*lacZ58*(M15), Δ*phoA8*, *glnX44*(AS), *deoR481*, *rfbC1*, *gyrA96*(Nal R), *recA1*, *endA1*, *thiE1* and *hsdR17* | Agilent Technologies, USA |
| *E. coli* BL21 (DE3) | $F^{-}$ *ompT gal dcm lon* $hsdS_B(r_B^{-}m_B^{-})$ λ(DE3 [*lacI lacUV5-T7p07 ind1 sam7 nin5*]) $[malB^{+}]_{K-12}(\lambda^{S})$ | Agilent Technologies, USA |
| *M. bovis* BCG | Vaccine strain against tuberculosis | A kind gift from Dr. Ramandeep Singh |
| pProEx-HTa | *E. coli* Expression vector with N-terminal 6xHis-Tag, $Amp^R$ | Invitrogen Inc., USA |
| pProEx-HTa::*phoR* | *phoR* gene encoding catalytic C-terminal domain (684-1458 bp), $Amp^R$ | (30) |
| pProEx-HTb::*gfp-phoR* | *phoR* gene encoding catalytic C-terminal domain (684-1458 bp) fused to GFP, $Amp^R$ | (57) |
| pET28a::*devR* | *devR* gene encoding DevR protein (1-654 bp), $Kan^R$ | (16) |
| pYA1556 | Full length PknK-pET SUMO, $Kan^R$ | (36) |
| pYA1659 | Full length PknK-K55M pET SUMO, $Kan^R$ | (39) |
